# Connectivity profile of thalamic deep brain stimulation to effectively treat essential tremor

**DOI:** 10.1101/575209

**Authors:** Bassam Al-Fatly, Siobhan Ewert, Dorothee Kübler, Daniel Kroneberg, Andreas Horn, Andrea A. Kühn

## Abstract

Essential tremor is the most prevalent movement disorder and is often refractory to medical treatment. Deep brain stimulation offers a therapeutic approach that can efficiently control tremor symptoms. Several deep brain stimulation targets (ventral intermediate nucleus, zona incerta, posterior subthalamic area) have been discussed for tremor treatment. Effective deep brain stimulation therapy for tremor critically involves optimal targeting to modulate the tremor network. This could potentially become more robust and precise by using state-of-the-art brain connectivity measurements. In the current study, we utilized two normative brain connectomes (structural and functional) to show the pattern of effective deep brain stimulation electrode connectivity in 36 essential tremor patients. Our structural and functional connectivity models were significantly predictive of post-operative tremor improvement in out-of-sample data (*p* < 0.001 for both structural and functional leave-one-out cross-validation). Additionally, we segregated the somatotopic brain network based on head and hand tremor scores. These resulted in segregations that mapped onto the well-known somatotopic maps of both motor cortex and cerebellum. Crucially, this shows that slightly distinct networks need to be modulated to ameliorate head vs. hand tremor and that those networks could be identified based on somatotopic zones in motor cortex and cerebellum.

Finally, we propose a multi-modal connectomic deep brain stimulation sweet spot that may serve as a reference to enhance clinical care, in the future. This spot resided in the posterior subthalamic area, encroaching on the inferior borders of ventral intermediate nucleus and sensory thalamus. Our results underscore the importance of integrating brain connectivity in optimizing deep brain stimulation targeting for essential tremor.

## Introduction

Essential tremor (ET) is the most common movement disorder that is encountered in clinical practice (Deuschl, 2000a). A satisfactory pharmacotherapeutic treatment is difficult if impossible to attain in 25-55% of ET cases (Flora *et al*., 2010). Therefore, deep brain stimulation (DBS) has been accepted as an efficacious alternative to control medication-refractory tremor symptoms.

To date, multiple DBS targets have been proposed to effectively treat ET (Deuschl *et al*., 2011). Targeting the VIM nucleus was regarded as a historical gold-standard since the beginnings of modern-day DBS (Benabid *et al*., 1991). Increasingly, the ventrally adjacent white matter has been proposed to lead to superior effects (Hamel *et al*., 2007; Sandvik *et al*., 2012; Eisinger *et al*., 2018). This target has been referred to as the posterior subthalamic area (PSA). Thus, the optimal treatment coordinates are still a matter of debate.

Pathophysiological evidence has accumulated that a cerebello-thalamo-cortical tremor network plays a crucial role in mediating abnormal oscillatory tremor activity and its modulation is related to the therapeutic effects of DBS (Schnitzler *et al*., 2009, Raethjen and Deuschl, 2012). The cortical and subcortical nodes constituting the proposed network have been described with fMRI and MEG (Sharifi *et al*., 2014, Schnitzler *et al*., 2009). In light of such a network-based mechanism, strong connectivity between DBS electrodes and network *tremor nodes* should lead to effective treatment response. This approach has been followed in individual cases by Coenen and colleagues who proposed DTI-based targeting in tremor patients focusing on the connectivity between the cerebellum and the thalamus (Coenen *et al*., 2011a, 2011b, 2017). Recently, a different approach has been proposed to use whole brain connectivity patterns to predict clinical outcome after DBS. This was first demonstrated in Parkinson Disease across cohorts, and improvement scores could be predicted across DBS centers and surgeons (Horn *et al*., 2017b, 2017a). In case of ET, few studies addressed the relationship between DBS connectivity and clinical outcome and so far, none has actually used brain connectivity to predict the DBS effects in out-of-sample data (Pouratian *et al*., 2011; Gibson *et al*., 2016; Akram *et al*., 2018; Middlebrooks *et al*., 2018).

Here, we aimed at constructing a “therapeutic network” model for DBS in ET. Following the concept of (Horn *et al*. 2017b), we postulated that similarity to this connectivity fingerprint could linearly predict clinical outcome in ET patients. We traced DBS-electrode connectivity to other brain regions using high resolution normative connectomes (functional and structural) as surrogate neuroimaging models in a data-driven fashion. We validated the resulting optimal connectivity fingerprints by predicting individual tremor improvements in a leave-one-out design. In a further step, we used DBS connectivity to investigate somatotopic treatment effects. Specifically, we analyzed how tremor improvement of hand and head could be associated with segregated DBS connectivity maps. Finally, we condensed findings to define an optimal surgical target for ET, which is made publicly available in form of a probabilistic atlas dataset.

## Materials and methods

### Patients: demographic and clinical details

Thirty-six patients underwent DBS (72 DBS electrodes) for severe, medically intractable ET (13 female) were retrospectively included in the current study (mean age = 74.3 ± 11.9 years). Diagnosis of ET followed the consensus criteria proposed in 1998 (Deuschl *et al*. 1998). Patients with bilateral symmetric postural or kinetic tremor of the upper limb with the possibility of additional head tremor, were included as ET cases. Any isolated voice, chin, tongue or leg tremor patients were excluded. Additionally, patients with dystonic, neuropathic, orthostatic, physiological or psychological tremor were excluded. Patients had a mean disease duration of 24.33 ± 14.99 years before DBS surgery. All patients received bilateral DBS implants in Charité–Universitätsmedizin, Berlin for the period between 2001 and 2017 (see Table 1 for clinical and demographic information and supplementary table S1 for individual patient clinical characteristics). All implanted DBS electrodes were Medtronic 3387 (except for three patients in which two were implanted with Boston Scientific Vercice Directed and one with St Jude Medical). Preoperative MRI was used to define VIM/zona incerta DBS targets. Microelectrode recordings and test stimulation were utilized intraoperatively to guide DBS lead placement. Correct lead placement was confirmed by postoperative imaging using LEAD-DBS to localize DBS electrodes in standard MNI space (Fig. 1). Percent improvement in the Fahn-Tolosa-Marin (FTM) tremor score served as an index of clinical outcome (Fahn *et al*., 1988). FTM scores before (baseline) and at least 3 months after electrode implantation have been obtained from archival video material. All videos have been rated by three clinicians experienced in movement disorders. Each clinician (BA, DKu and DKo) rated separate part of the cohort (so no video was rated twice) and was blinded to the timing of the video (preoperative vs postoperative). Postoperative FTM scores indicate tremor severity during the chronic DBS ON condition. Upper limb subscores contralateral to DBS electrodes were summed up and used in the calculation of the main clinical outcome quantifying therapeutic effect. Upper limb subscore comprised the following items: rest tremor, postural tremor, action tremor, drawing of Archimedes spiral and repeated letter L writing (modified FTM score). For somatotopy related analyses, bilateral upper limb subscores and head scores were used. The head score consisted of the sum of head, face, tongue, speech and voice related subscores. All patients showed a reduction in FTM score of at least ∼ 27% with a mean decrease of 22.4 ± 9.9 points of the average total FTM score (from 33.3 ± 9.6 at baseline to 10.9 ± 5.5 with chronic DBS). The average postoperative time at which patients were assessed for postoperative FTM scoring was 12 ± 9.86 months.

**Figure 1:**
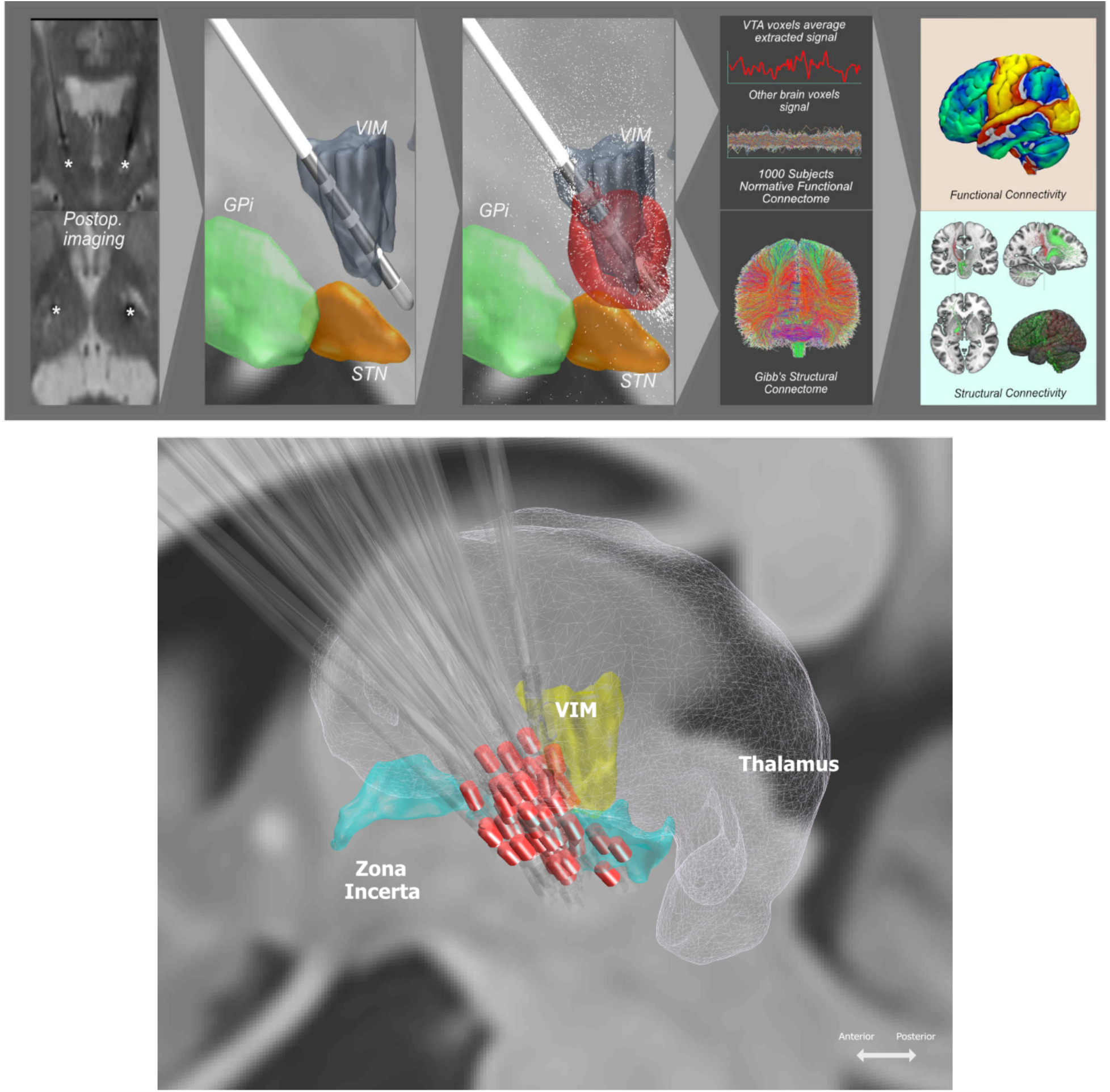
Upper panel: Methodological pipeline of data analysis: A. DBS leads were localized using Lead-DBS software. B. 3D reconstruction of the DBS lead in standard space. C. Modeling volume of brain tissue electrically activated by the active electrode contact (VTA, red). Estimating functional (D) and structural (E) connectivity metrics using normative connectomes. Connectivity was calculated between the volume of tissue activated as a seed and the rest of the brain. F. Building predictive models by correlating the connectivity metrics to clinical improvement. Lower panel: Deep brain stimulation electrode localizations in standard space. Red colored marks active contacts. All DBS leads shown on the left side after flipping right sided electrodes.

**Table 1:** Cohort demographics and clinical data.

| Criteria |  |
| --- | --- |
| Age, years | 74.3 ± 11.9 |
| Age at diagnosis, years | 44.9 ± 18.4 |
| Disease duration, years | 24.3 ± 14.9 |
| Male sex, (%) | 23 (72) |
| Baseline total FTM score | 33.3 ± 9.6 |
| Postoperative total FTM score | 10.9 ± 5.5 |
| Total FTM improvement (%) | 65.1 ± 18.4 |
| Baseline contralateral UL tremor score | 13.4 ± 4.3 |
| Postoperative contralateral UL tremor score | 4.6 ± 2.9 |
| Contralateral UL tremor improvement (%) | 63.4 ± 22.9 |
| Baseline head tremor score | 3.8 ± 2.8 |
| Postoperative head tremor score | 1.0 ± 1.7 |
| Head tremor improvement (%) | 80.8 ± 29.5 |

UL: upper limb, FTM: Fahn-Tolosa-Marin tremor score.
Data are presented in mean ± SD.
Absolute tremor score were reported at baseline and postoperative time points while tremor improvement were reported in percentage.

The study was approved by the local ethics committee of the Charité University Medicine - Berlin.

### DBS electrode localizations

Preoperative MRI as well as postoperative MRI or CT were obtained in all patients. DBS electrodes were localized using Lead-DBS software (Horn & Kühn 2017; www.lead-dbs.org) following the enhanced methodology described in (Horn & Li *et al*. 2018 NeuroImage). Briefly, preoperative and postoperative patients’ images were linearly co-registered using Advanced Normalization Tools (ANTs, Avants *et al*., 2009; http://stnava.github.io/ANTs/) and manually refined when necessary.

Pre- and postoperative images were then normalized into ICBM 2009b NLIN asymmetric space using the symmetric diffeomorphic image registration approach implemented in ANTs (Avants *et al*., 2009; http://stnava.github.io/ANTs/). Electrodes were then localized and volumes of tissue activated (VTA) modeled using Lead-DBS based on patient-specific stimulation parameters.

### Functional and Structural Connectivity Estimation

Using VTAs as seed regions, functional and structural connectivity estimates were computed using pipelines implemented in Lead-DBS. Two normative connectomes were used: First, a structural connectome (Horn *et al*., 2014; Horn, 2015) which consists of high density normative fibertracts based on 20 subjects. Diffusion data were collected using single-shot spin-echo planar imaging sequence (TR = 10,000 ms, TE = 94 ms, 2 × 2 × 2 mm^3^, 69 slices). Global fiber-tracking was performed using Gibb’s tracking method (Reisert *et al*., 2011) (for more methodological details, see Horn and Blankenburg (2016)). Structural connectivity was estimated by extracting tracts passing through VTA seeds and calculating the fiber counts in a voxel-wise manner across the whole brain. Second, a functional connectome which was defined on 1000 healthy subjects resting-state fMRI scans (Yeo *et al*., 2011; https://dataverse.harvard.edu/dataverse/GSP) and is based on data of the Brain Genomics Superstruct Project. Data were collected with 3T Siemens (Erlangen, Germany) MRI and the resting state BOLD processed with signal regression and application of spatial smoothing kernel of 6mm at full-width at half maximum (Yeo *et al*., 2011). For the purpose of the current study, connectivity estimates were performed for each of the 72 VTAs (36 bilateral implants) after nonlinearly flipping right sided VTA to the left side using Lead-DBS.

### Models of optimal connectivity

Following the concept described in (Horn et al., 2017b), clinical improvements in the contralateral upper limb were correlated with structural and functional connectivity from the VTA (while these were accumulated on the left side of the brain) to each brain voxel across electrodes. This process resulted in R-maps that carry Spearman’s rank-correlation coefficients for each voxel. The maps fulfill two concepts. First, they denote to which areas connectivity is associated with beneficial outcome. Second, their spatial distribution describes an optimal connectivity profile of DBS electrodes for ET (Horn *et al*., 2017b).

Thus, to make predictions, each VTA-derived structural or functional connectivity pattern was then tested for spatial similarity with this optimal connectivity model. Specifically, similarity between each VTA’s connectivity profile and the “optimal” connectivity profile (as defined by the R-map) was calculated using spatial correlation. The resulting similarity index estimates “how optimal” each connectivity profile was and was used to explain clinical improvement in a linear regression model. To cross-validate the model, we correlated aforementioned predicted and empirical individual upper limb tremor improvements in a leave-one-out design. Furthermore, we calculated discriminative fibertracts following the approach introduced recently by Baldermann and colleagues (Baldermann et al. 2019). Briefly, fibertracts connected to VTAs across cohort were isolated from the normative group connectome. In a mass-univariate analysis, for each fibertract, a two-sample t-test was performed between improvement scores of VTAs connected versus improvement of non-connected VTAs and fibers were labelled according to this t-score. The resulting positive t-score streamlines represent fibertracts that may discriminate between poor and good responders. Again, this analysis was performed across the left-sided accumulated VTAs using contralateral upper limb improvement subscores. This analysis was used to confirm the main analysis using a slightly different statistical concept.

#### Prospective Case Validation

We pre-operatively scanned one patient with diffusion weighted imaging (see supplementary methods for scan parameters) to investigate the validity of our model in predicting patient improvement using patient-specific tractography. The patient received an unilateral implant on the left (Abbott’s St. Jude Medical Infinity model) for treatment of refractory ET affecting the upper limbs. The VTA was modeled with the same pipeline as the main patients cohort. Patient-specific diffusion weighted imaging (dMRI) data was then used to calculate fiber streamlines seeding from the modeled left-VTA. The resulting connectivity profile was then fed into the structural predictive model created on the main cohort (using the normative connectome). Patient’s empirical right upper limb FTM score was calculated pre- and post-operatively following the same methodological description as in the main cohort.

#### Side-effects Related Connectivity Profile

Connectivity seeding from electrodes associated with DBS-related side effects were also calculated in a subgroup of patients in which information about side effects were available using the same functional connectome (Yeo et al. 2011). We then compared the resulting connectivity to a sample of control patients DBS-induced side-effects could be excluded. To do so, mass-univariate voxel-wise two-sample t-tests were calculated between connectivity strengths seeding from VTAs associated with gait ataxia or dysarthria and that of control patients. Connectivity difference images were then masked by significant p-values (<0.05, uncorrected) and presented as positive t-scores images.

### Deriving Somatotopic Maps

In a further step, we segregated somatotopic maps informed by optimal functional connectivity models based on upper limb (hand) and head tremor improvements. Since head tremor is an axial feature modulated by both left and right VTAs, those were combined in this analysis. Hence, bilateral VTAs were used to estimate functional somatotopic maps (i.e. connectivity was estimated seeding from both VTAs). The resulting connectivity maps were correlated with either summed bilateral hand scores or head scores. The resulting R-maps were overlaid on the cerebellum and primary motor cortex to investigate somatotopy.

### Defining an optimal DBS target

As a final step, we applied our optimal predictive structural and functional models to define an “optimal” DBS target. We masked our functional and structural R-maps to include only voxels in the cortical and cerebellar regions. This was done since otherwise the design would have been recursive (with subcortical information already present in the R-maps). The subcortical region with maximal connectivity to those R-maps was determined using Lead-DBS. The resulting connectivity maps were then overlapped to show where exactly they converged. This spot is characterized by optimal functional and structural brain connectivity for maximal therapeutic outcome.

### Data availability

Patients dataset are not publicly available due to data privacy restrictions but can be made available from the corresponding author upon reasonable request. All code used in the present manuscript is available within Lead-DBS software (https://github.com/leaddbs/leaddbs).

## Results

In total, 72 DBS electrodes were included in the analyses. Connectivity based R-maps highlighted positively predictive voxels in multiple regions (Fig. 2) such as paracentral gyrus (M1 and sensory cortex), visual cortices (V1 and V2), superior temporal gyrus, and superior and inferior cerebellar lobules. Additionally, functional connectivity to part of the premotor cortex and supplementary motor area was associated with beneficial DBS outcome. On the other hand, structural optimal connectivity outlined additional regions such as superior parietal lobule and precuneus. Except those, the beneficial functional and structural connectivity profiles were largely congruent.

**Figure 2:**
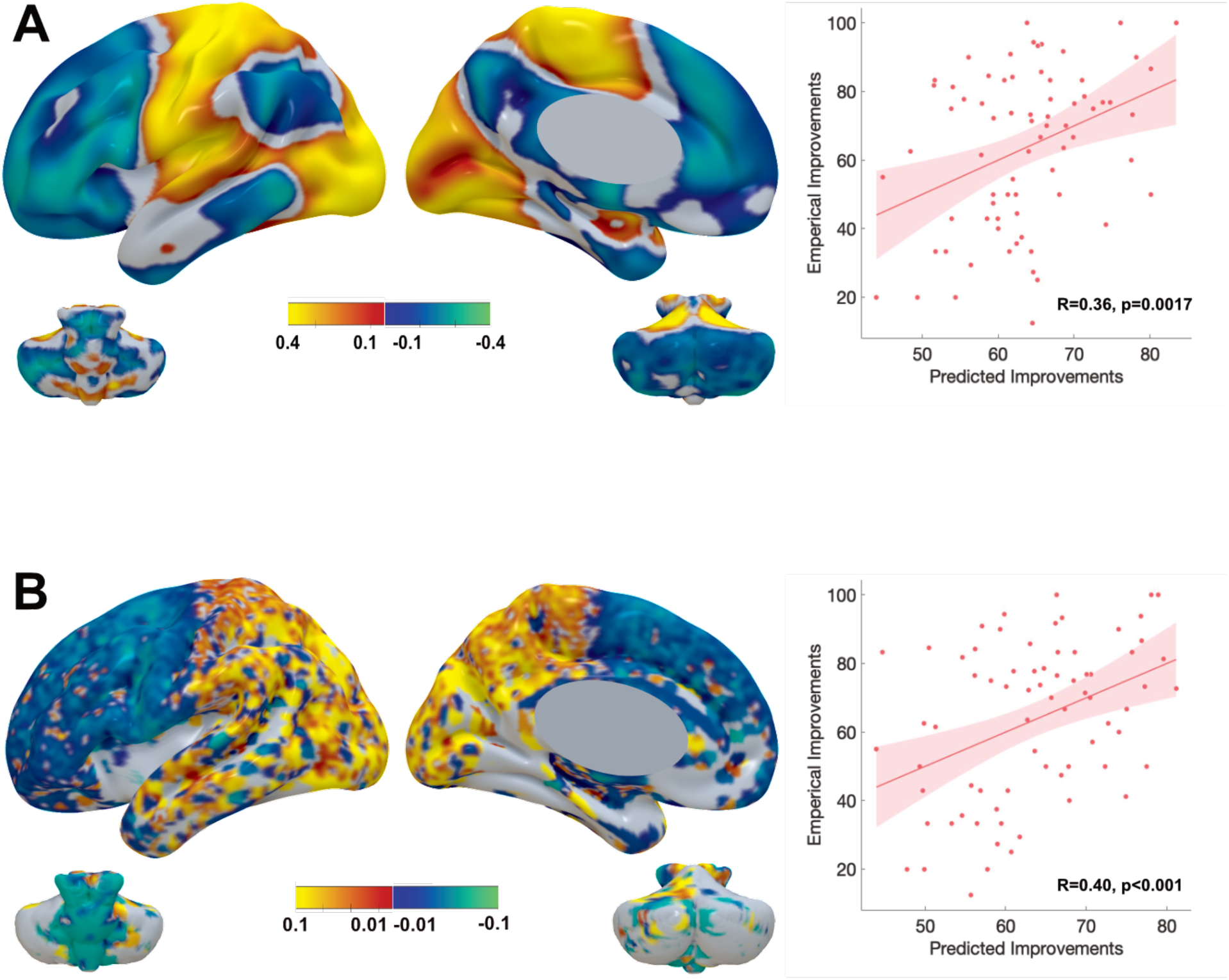
A. Functional connectivity predictive of clinical improvement. Voxel topology predictive of DBS outcome generated using a high-definition functional connectome. The scatter plot demonstrates the correlation between predicted improvement (based on similarity between predictive functional connectivity profiles and functional connectivity profiles seeding from each VTA) and original clinical improvement scores of sixty-six upper limbs in a leave-one out design (R = 0.36, *p* = 0.002). B. Topological distribution of structural connectivity predictive of DBS related improvement. Connectivity generated using normative structural connectome. Voxels containing streamline counts that were positively correlated to clinical improvement are shown (bottom left). Result of leave-one out cross validation (R = 0.40, p < 0.001) is shown in the scatter plot.

Functional connectivity profiles could explain 16.4% of the variance in DBS outcome (R = 0.41, *p* < 0.001), while structural connectivity profile could explain 25% of the variance in DBS outcome (R = 0.50, *p* < 10^-5^). In a leave-one-out cross-validation, both structural (R = 0.40, *p* < 0.001) and functional connectivity (R = 0.36, *p* = 0.0017) remained significant predictors of individual clinical improvement. On average, predicted tremor improvements deviated from empirical improvements by 17.98 ± 10.73 % for structural and 18.09 ± 11.22 % for functional connectivity. As a proof of concept, similarity between VTA-seed connectivity in one modality and the R-map model of the other was also significantly predictive of tremor improvement (functional VTA-seed connectivity explained by structural model R = 0.41, *p* < 0.001; structural connectivity explained by functional model R = 0.33, *p* = 0.005). This may further illustrate similarities between optimal functional and structural connectivity maps. While our main analysis focused on improvements of hand-tremor scores, we repeated the main analysis for improvements of full tremor scores which led to near identical results (see Fig. S1).

Structural DBS connectivity showed voxel clusters intersecting with a DBS target commonly used in ET treatment (Papavassiliou *et al*., 2004) and with the cerebello-thalamo-cortical tract (Fig. 3). The cluster extended from the M1 cortex down to the thalamic-subthalamic region. Discriminative fibertract analysis delineated a well-defined tract connecting M1 and cerebellum (Fig. 5), passing through the motor thalamus. Crucially, based on our results, this tract represented the part of the cerebello-thalamo-cortical pathway that was associated with optimal improvement.

**Figure 3:**
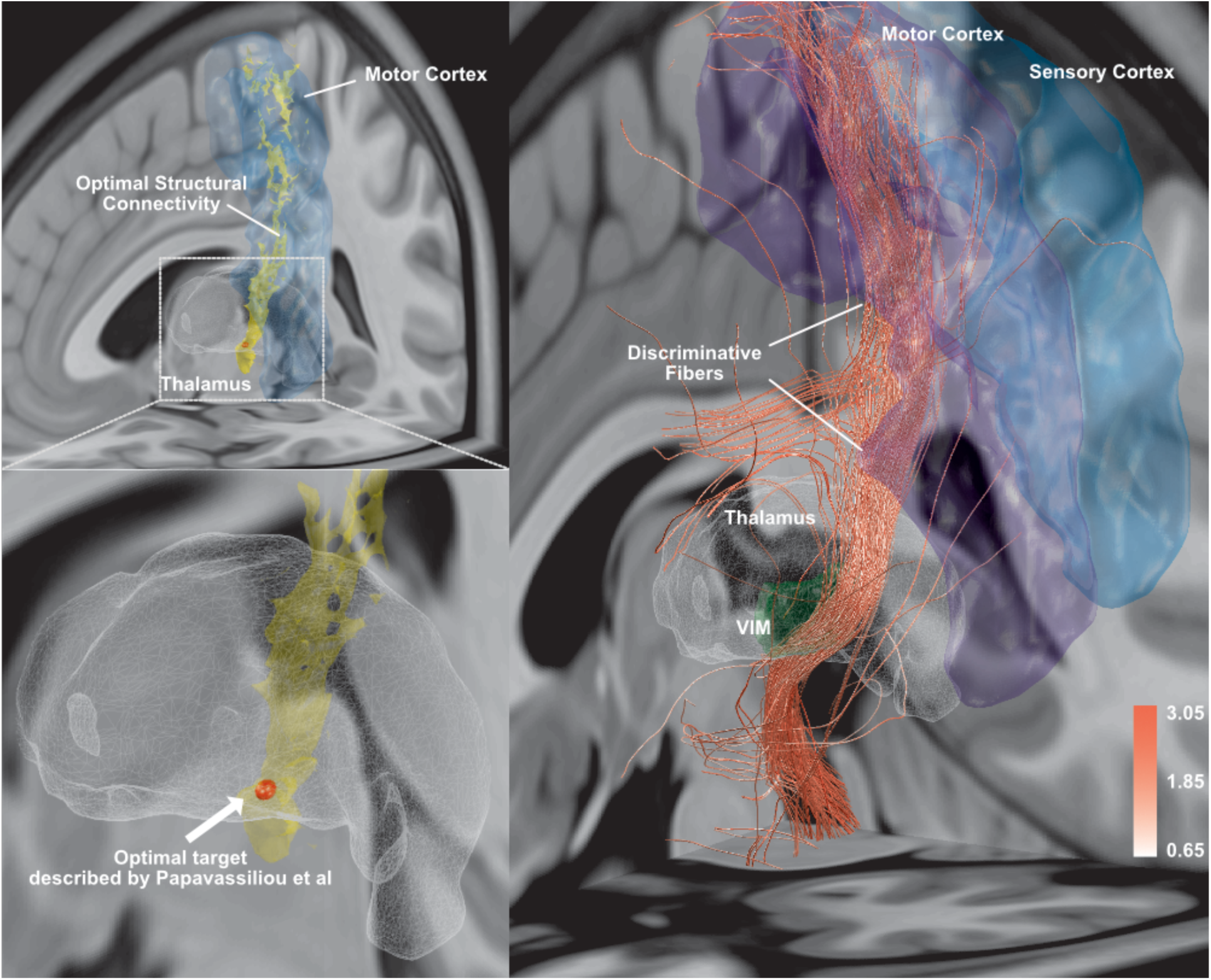
Overlap of predictive voxels in structural connectivity model with literature-based DBS. Voxels extend from the area of M1 to the thalamic-subthalamic region. Discriminative fibertracts predictive of DBS outcome were statistically delineated and correspond well to the cerebello-thalamo-cortical pathway (right panel). Of note, tracts crossing the corpus callosum as well as non-decussating tracts toward the cerebellum are likely false-positive tracts commonly observed using diffusion-based tractrography. Color-bar represents t-scores of discriminative streamlines.

**Figure 4:**
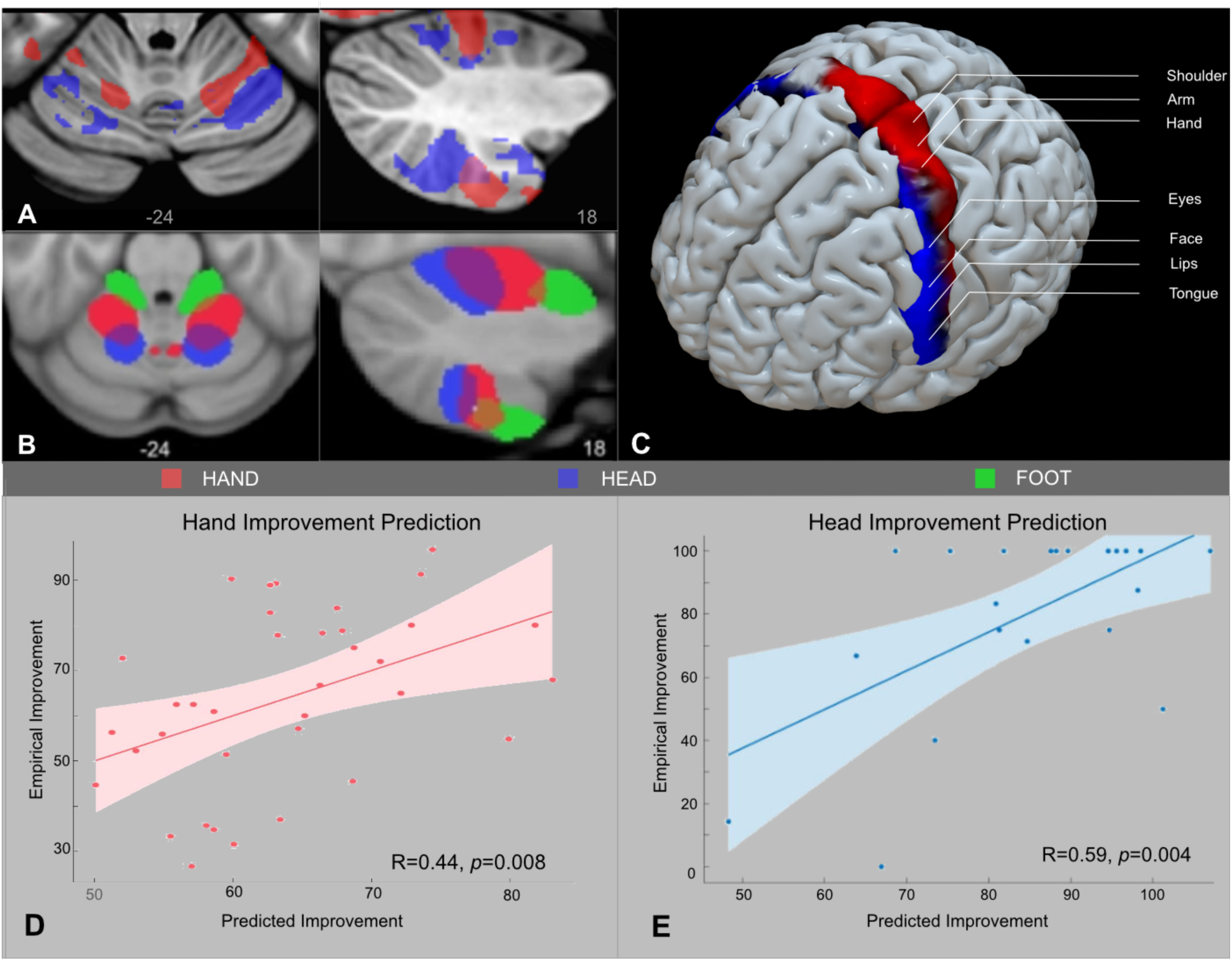
A. Results from current study and B. a previous resting state fMRI study performed in healthy subjects (Yeo *et al*., 2011). Regions of hand and head tremor score in the cerebellar gray matter conform to formerly depicted regions for hand and tongue somatotopic regions of the cerebellum (Buckner *et al*., 2011). C. Motor cortex distribution of regions associated with hand and head tremor score correspond to the well-known homuncular structure of M1. D. Prediction of hand tremor improvement score (36 patients) using DBS connectivity to combined cerebellar and motor hand regions. E. Prediction of head tremor improvement score (22 patients) using DBS connectivity to combined cerebellar and motor regions.

**Figure 5:**
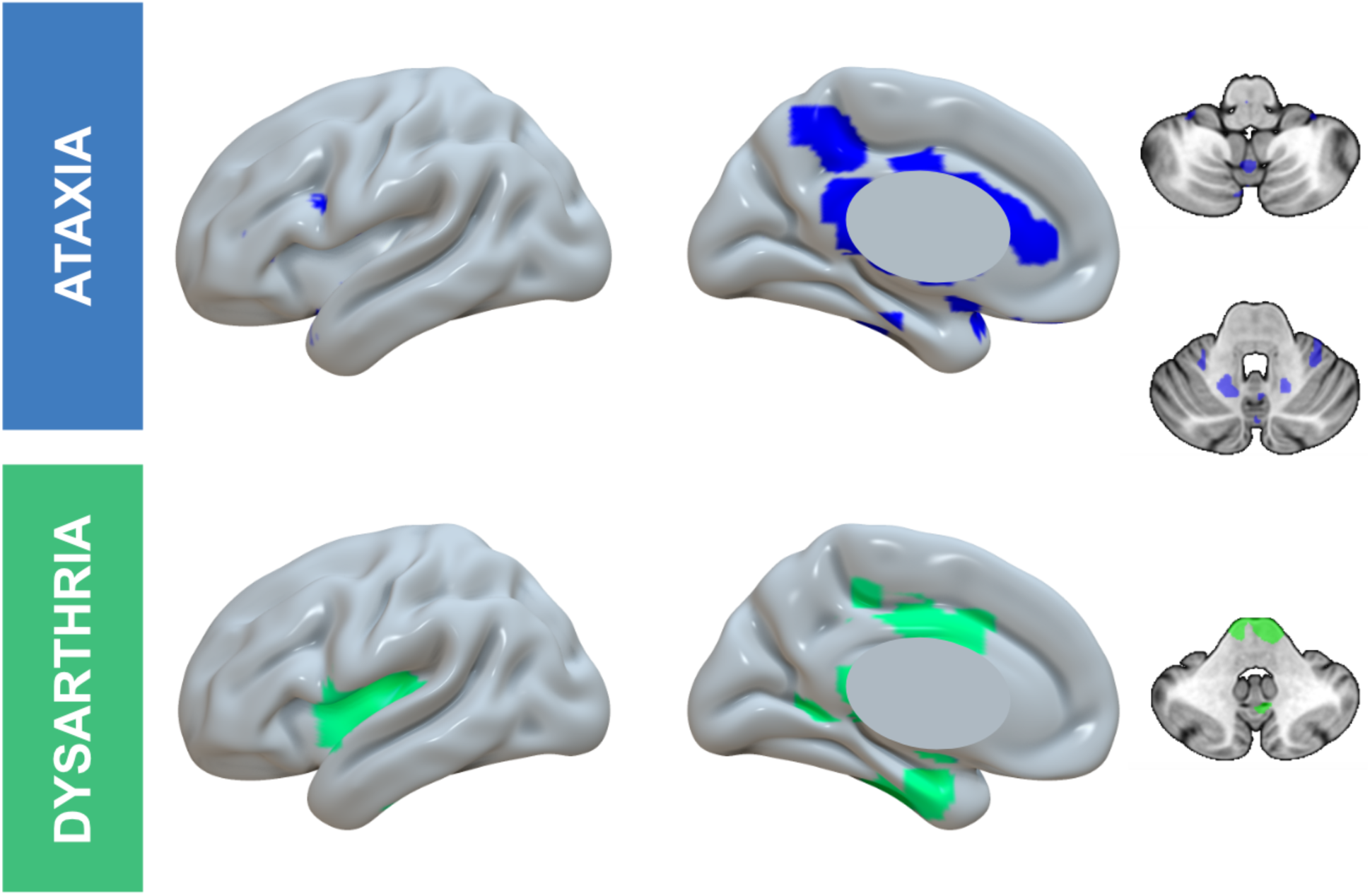
Connectivity patterns associated with gait ataxia and dysarthria as representative VIM DBS induced side-effects. Regions highlighted in the figure were associated with occurrence of these commonly encountered side-effects (p < 0.05, uncorrected).

Beneficial structural connectivity (based on normative connectome) successfully predicted the magnitude of tremor improvement in a single prospective patient (empirical clinical improvement 61%, predicted clinical improvement 72%). This prediction was performed using patient-specific structural connectivity (Fig. S2).

Next, we aimed at defining functional connectivity maps that could explain therapeutic response in different body parts (hand vs. head tremor, Fig. 4). Of note, only 22 patients were included in the functional connectivity model of head tremor since the symptom was not present in the remaining 11. All patients responded well to head-tremor at baseline, thus a subanalysis comparing good vs. bad responders was not possible. The topology of M1 and cerebellar voxels predictive of hand and head tremor improvement followed the known homuncular organization of M1 and somatotopy of the cerebellum (Buckner *et al*., 2011). Furthermore, connectivity to these somatotopy-specific sub-regions of the cerebellum and M1 could explain improvement of hand (R = 0.44, *p* = 0.008), and head tremor (R = 0.59, p = 0.004), respectively.

Additionally, we investigated functional connectivity patterns that could differentiate patients with DBS-related side-effects (namely gait ataxia and dysarthria) from control patients. Our analysis revealed side-effect specific clusters. Interestingly, these cortical and cerebellar clusters overlapped minimally with voxels positively correlated with optimal DBS outcome. Of note, these results are not corrected for multiple comparisons and should be interpreted with caution.

Our final goal was to define a clinically relevant surgical target that maximizes beneficial connectivity within the thalamo-subthalamic area. In order to obtain such target, we seeded back from cortical voxels in our structural and functional R-maps (using their entries as a weighted connectivity seeds in Lead-DBS). Only cortical voxels were included to avoid confusion with already highlighted voxels in the sub-cortex. The resulting functional and structural connectivity patterns converged at the inferoposterior border of the VIM and extended inferiorly and posteriorly to overlap with the dorsal part of the zona incerta (Fig. 6).

**Figure 6:**
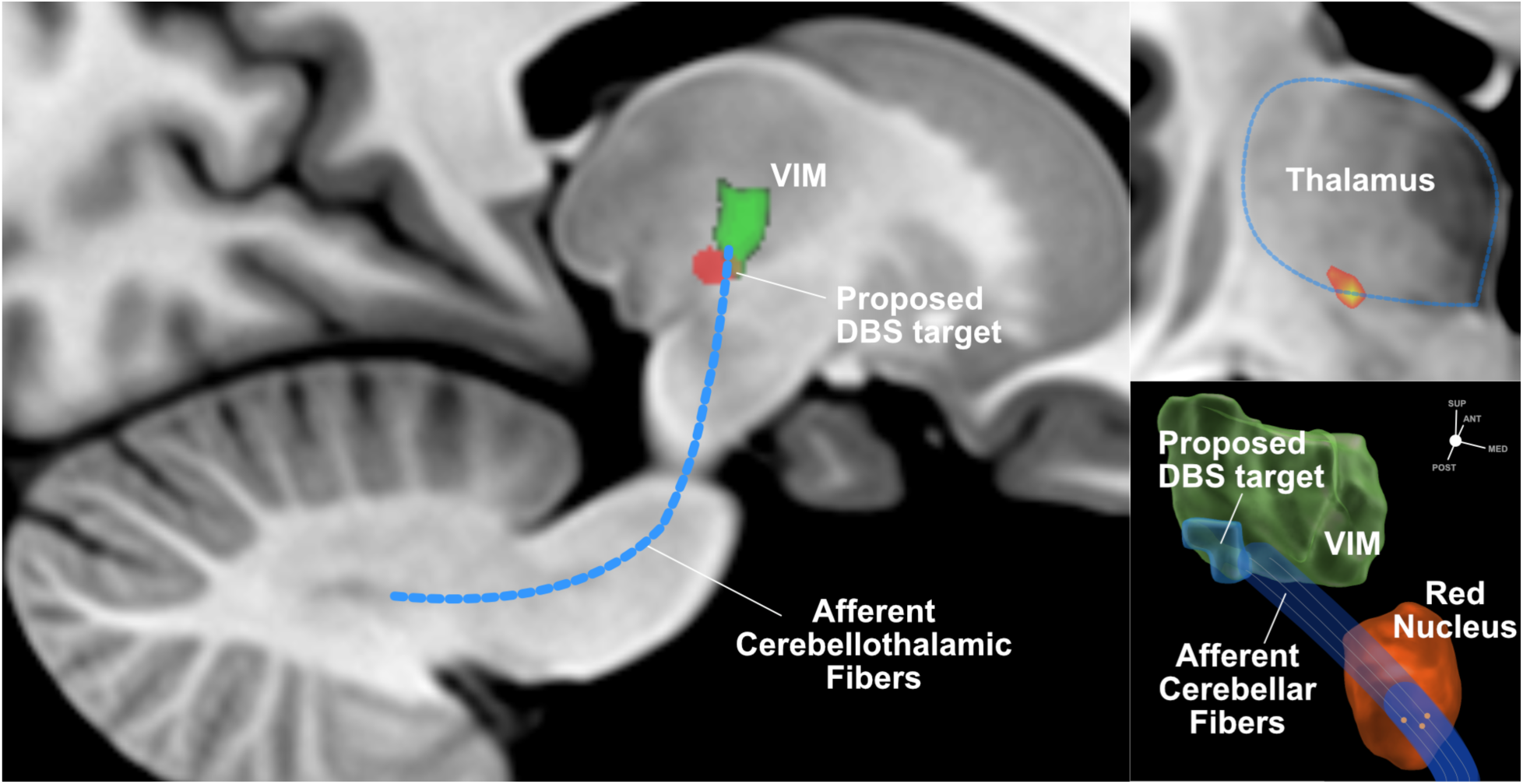
Connectivity-defined optimal location for DBS placement in essential tremor patients. Left: Sagittal view of MNI152 space showing VIM (red) and DBS target (green) derived from beneficial connectivity. The location of the proposed target is directly adjacent to the VIM (posteroinferiorly) in a subthalamic region where afferent cerebellothalamic fibers approach the VIM nucleus. Right upper:. Coronal view showing the spatial relation between the connectivity-based DBS target and the thalamus (ventrolateral location). Right lower: 3D schematic reconstruction of VIM (green). and Red Nucleus (red) showing the location of connectivity-based DBS target (blue) and its intersection with cerebello-thalamic fibers.

## Discussion

We demonstrated that optimal tremor reduction with DBS is significantly correlated with a specific pattern of functional and structural connectivity including sensorimotor areas and cerebellum. Importantly, the connectivity fingerprint of brain tissue activated by DBS can predict tremor improvement in out-of-sample data. Our models of optimal “therapeutic connectivity” largely overlap with brain regions that were linked to ET pathophysiology, before. More importantly, we demonstrated that tremor in distinct body parts is optimally ameliorated by modulating a specific network that includes somatotopic regions of both M1 and the cerebellum.

Finally, we defined an “optimal” DBS target that maximizes beneficial functional and structural connectivity.

### The tremor network and pattern of beneficial DBS connectivity

The mechanism of tremor generation has been attributed to multiple central oscillators (Schnitzler *et al*., 2009) that are synchronized in a tremor specific frequency (Marsden *et al*., 2000; Hellwig *et al*., 2001) and distributed across nodes of the cerebello-thalamo-cortical pathway. It has been thought that the cerebellum drives tremorogenic oscillations (Deuschl, 2000b). However, several studies unveiled the involvement of cortical (sensorimotor, supplementary motor and premotor cortices) and subcortical (thalamus) nodes in tremor generation (McAuley, 2000; Pinto *et al*., 2003; Schnitzler *et al*., 2009; Helmich *et al*., 2013). Theoretically, interference with any of these cerebello-thalamo-cortical nodes should suppress tremor oscillation. The thalamic (VIM) nucleus which receives most of the cerebellar afferent fibers (Asanuma *et al*., 1983) has been of much interest in tremor research (Pedrosa *et al*., 2012; Basha *et al*., 2014; Fang *et al*., 2016; Milosevic *et al*., 2018). The VIM also projects to the aforementioned tremor-related motor areas (McFarland and Haber, 2002; Haber and Calzavara, 2009). This property gives it a central position in the cerebello-thalamo-cortical tremor pathway. Historically, it was considered an excellent target for lesioning surgery (thalamotomy) yielding a satisfactory outcome of tremor control (Deuschl *et al*., 2011). Later, DBS surgery started to replace thalamotomy in the majority of cases, given its reversible and adjustable stimulation (Tasker, 1998). Nonetheless, clear visualization of the VIM region with conventional MRI is difficult even in contemporary DBS surgery with modern imaging protocols (Yamada *et al*., 2010), this is why connectivity has already been used to target VIM-DBS surgeries (Anderson *et al*., 2011; Coenen *et al*., 2014).

This said, the optimal DBS target has to have tight functional and structural connectivity to the tremorogenic nodes in order to remotely modulate the nuisance tremor oscillations. Our results showed a connectivity pattern which agrees with this concept. Both structural and functional connectivity demonstrated areas in the pre- and postcentral gyri in addition to the superior and inferior cerebellar lobules. This is in line with the results of most studies that showed tremor related alterations of the sensorimotor and cerebellar areas (Colebatch, 1990; Jenkins *et al*., 1993; Wills *et al*., 1995; Czarnecki *et al*., 2011; Fang *et al*., 2013; Mueller *et al*., 2017). Additionally, target connectivity to the aforementioned areas was associated with tremor improvement in VIM-DBS and ablative (thalamotomy) surgeries (Klein *et al*., 2012; Gibson *et al*., 2016; Akram *et al*., 2018; Middlebrooks *et al*., 2018; Tuleasca *et al*., 2018a).

Other regions that could potentially play a role based on present findings are primary and associative visual cortices. The importance of brain visual areas in tremor pathogenesis has been recently investigated by using a visual task of increasing difficulty to illustrate the impact of visuospatial network in tremor augmentation (Archer *et al*., 2018). Furthermore, recent series of investigations suggested that structural and functional changes of the visual cortex could be a preoperative predictor of optimum tremor outcome after ablative radiosurgery (Tuleasca *et al*., 2017, 2018b, 2018c).

### Somatotopic organization of beneficial DBS connectivity

Finely tuned DBS targeting with respect to the somatotopy of body regions has been considered in dystonia patients (Vayssiere *et al*., 2004). We leveraged the nature of anatomical somatotopic distributions in order to explain how DBS related connectivity profile could vary accordingly. Our results demonstrated two distinct connectivity profiles corresponding to hand and head in M1 and cerebellar regions. Crucially, these areas corresponded to formerly determined hand and tongue brain regions in the human M1 homunculus (Penfield and Boldrey, 1937) and cerebellum (Buckner *et al*., 2011). Furthermore, they predicted DBS tremor reduction in their respective body regions. Our finding supports the utility of hand and head tremor driven connectivity profiles in guiding DBS targeting, which could be an important future step for further refinement of DBS treatment of focal motor symptoms. Head tremor is the second most common body distribution of tremor symptoms encountered in ET patients that is highly disabling beside the predominant upper limbs tremor (Hoskovcová *et al*., 2013; Bhatia *et al*., 2018). Correspondingly, controlling head tremor has been an outcome issue in many patients undergoing DBS surgery (Obwegeser *et al*., 2000; Putzke, 2005). Our results may pave the way for personalized DBS targeting that is dependent on the tremor symptoms each patient may have. It is even conceivable to scan patients in the fMRI while they perform (imaginary) tasks involving hand and head to identify their specific somatotopic organization of M1 and the cerebellum. These regions could then be used in single patients to define the tremor target optimally corresponding to their symptomatology.

### Connectivity-derived predictive models

The beneficial connectivity profiles that were estimated in the present work were built using a completely data-driven design. This means that these profile maps can be interpreted as an answer to where in the brain connectivity may explain most of the variance in clinical improvement. The concept of using connectivity patterns to predict functional capacity and clinical symptoms has been a central dogma in contemporary studies (Beaty *et al*., 2018; Cao *et al*., 2018). We relied on this concept in order to draw conclusions about the optimal connectivity fingerprint that will ensure the best outcome. Of note, connectivity associated with the emergence of side-effects involved inverse patterns of brain areas compared to beneficial DBS outcome. The cerebellar vermis was shown as a key region in ataxia related analysis, which is in accordance with previous results (Reich et al., 2016). Our models could significantly predict tremor improvement in out-of-sample data as well as in a single prospective patient using patient-specific dMRI data. Future work should focus on validating such connectivity fingerprints in a larger sample of prospective patients. Furthermore, the isolated discriminative tract emphasized the importance of targeting cerebello-thalamo-cortical pathways for determining DBS outcome (Coenen et al., 2014, Sammartino et al., 2016).

### A connectomic DBS target for Essential Tremor

The exact DBS target for optimal therapeutic benefit in ET is not yet entirely clear. Four main surgical targets have been suggested for essential tremor treatment. Located within the thalamus, the VIM nucleus has been regarded as the mainstay therapeutic target (Benabid *et al*., 1991; Pahwa *et al*., 2006; Zhang *et al*., 2010; Baizabal-Carvallo *et al*., 2014), while the other three targets within the subthalamic area (the PSA, which encompasses caudal zona incerta and the radiation prelemniscalis, and subthalamic nucleus) were the focus of other studies (Herzog *et al*., 2004; Plaha *et al*., 2008; Fytagoridis and Blomstedt, 2010). VIM DBS has proven to be an effective tremor target since the beginnings of modern-day DBS (Benabid *et al*., 1991; Deuschl *et al*., 2011). On the other hand, there is a growing evidence that DBS to the directly adjacent PSA is similarly effective (Plaha *et al*., 2004, 2011; Fytagoridis *et al*., 2012; Barbe *et al*., 2018). Deciding which target is optimal for tremor suppression is a critical step in stereotactic surgery. The results of the present study showed that the discussed targets may in fact be the same – fibers that pass along the red nucleus toward the thalamus and in doing so traverse through the PSA and zona incerta. Structural and functional connectivity maps converged in a region that impinge the inferior-thalamic border and extend to the PSA. Moreover, the proposed DBS spot is located ventrolateral to the thalamus, in an area medial to the internal capsule and directly inferior to the VIM and sensory thalamic nuclei, encroaching on their inferior borders. This area has been described in the literature as the entry of the afferent cerebellar fibers to the thalamus (particularly, the VIM nucleus; Gallay *et al*., 2008). Our results further imply the importance of the cerebellothalamic tremor pathway and encourage tract-based targeting for ET treatment (Sammartino *et al*., 2016; Fenoy and Schiess, 2018). Intriguingly, the identified spot is in accordance with a recently described optimal location for focused ultrasound thalamotomy in essential tremor treatment (Boutet *et al*., 2018) and with a previously published sweet spot (Papavassiliou *et al*., 2004).

## Limitations of the study

We used normative connectome data to estimate seed-based connectivity in individual patients. This concept has been introduced for studies in clinical domains such as stroke (Darby *et al*., 2018; Joutsa *et al*., 2018a, 2018b), DBS (Horn *et al*. 2017b, Fox *et al*., 2014) or TMS (Weigand *et al*., 2018) where patient-specific connectivity data is often lacking. Although these connectome atlases do not represent patient-specific connectivity, they in turn have the benefit of high signal-to-noise ratios. The functional connectome we used was defined on a high N (1000 subjects) and was acquired using specialized MR hardware (Yeo *et al*., 2011). In addition, the structural connectome was calculated using a modern approach that was best performer among 10 different tractography processing algorithms in an open competition (Fillard *et al*., 2011). Finally, this limitation should bias our results toward non-significance to predict out-of-sample data, but instead, the models proved highly robust in cross-validation.

Second, the retrospective design of our study poses a limitation. Needless to say, our exemplary attempt to validate the model on a single case scanned with patient-specific diffusion MRI should only be considered as an anecdotal evidence. Despite the good performance of our models in predicting individual outcome, a prospective multi-center study is needed to translate our results into clinical practice. Additionally, our side-effects connectivity analysis was based on small number of patients and did not involve a quantitative assessment of side-effects. As a consequence, results did not survive correction for multiple comparisons. Nevertheless, these results could be used to form hypotheses for further studies that may specifically address the connectivity fingerprints of VIM-DBS induced side-effects.

Third, inter-individual anatomical variability implies another challenge in predicting individual optimal DBS target using an optimal target from a group analysis. Nevertheless, our target was built on a connectome-based model which emphasizes the importance of targeting structural and functional connectivity between DBS electrode and regions delineated by the predictive models (specifically M1 and the cerebellum).

Lastly, our cohort assumed a single category of tremor syndromes, namely essential tremor. this could be of concern since other tremor syndromes equally benefit from DBS surgery (Kumar *et al*., 2003; Herzog *et al*., 2004; Foote *et al*., 2006; Mandat *et al*., 2010; Kilbane *et al*., 2015; Cury *et al*., 2017). For example, Parkinsonian tremor is successfully treated with subthalamic nucleus (Diamond *et al*., 2007) and VIM (Kumar *et al*., 2003) DBS. How connectivity patterns of effective DBS therapy could predict tremor reduction across different targets and tremor semiology remains to be established.

## Conclusion

We identified patterns of connectivity that allow to predict individual clinical outcomes of DBS in ET patients. More specifically, we introduced somatotopic connectivity maps that bear the potential of steering DBS targeting and programming toward patient-specific profiles with respect to the body distribution of symptoms. Finally, we estimated an “optimal” DBS target and set it into relationship to known ET-DBS targets. Our target is based on the convergence of beneficial functional and structural connectivity patterns and is available as a probabilistic, deformable atlas that we made openly available within the software Lead-DBS. Our results add to the ongoing effort of connectivity-based DBS targeting and foster the advance of connectomic surgery.

## Acknowledgements

We would like to thank Igor Ilinsky and Kristy-Kultas Ilinsky (University of Iowa) for their helpful comments on our results. We also thank Wolf-Julian Neumann (Charité – University Medicine Berlin) for his valuable advice and comments.

## References

Akram H, Dayal V, Mahlknecht P, Georgiev D, Hyam J, Foltynie T, et al. Connectivity derived thalamic segmentation in deep brain stimulation for tremor. NeuroImage Clin 2018, 18:130–142.

Anderson JS, Dhatt HS, Ferguson MA, Lopez-Larson M, Schrock LE, House PA, et al. Functional Connectivity Targeting for Deep Brain Stimulation in Essential Tremor. Am J Neuroradiol 2011, 32:1963–8.

Archer DB, Coombes SA, Chu WT, Chung JW, Burciu RG, Okun MS, et al. A widespread visually-sensitive functional network relates to symptoms in essential tremor. Brain 2018, 141:472–485.

Asanuma C, Thach WT, Jones EG. Brainstem and spinal projections of the deep cerebellar nuclei in the monkey, with observations on the brainstem projections of the dorsal column nuclei [review]. Brain Res Rev 1983, 5:299–322.

Avants B, Epstein C, Grossman M, Gee J. Symmetric diffeomorphic image registration with cross-correlation: Evaluating automated labeling of elderly and neurodegenerative brain. Med Image Anal 2008, 12:26–41.

Baizabal-Carvallo JF, Kagnoff MN, Jimenez-Shahed J, Fekete R, Jankovic J. The safety and efficacy of thalamic deep brain stimulation in essential tremor: 10 years and beyond. J Neurol Neurosurg Psychiatry 2014, 85:567–572.

Baldermann JC, Melzer C, Zapf A, Kohl S, Timmermann L, Tittgemeyer M, et al. Connectivity Profile Predictive of Effective Deep Brain Stimulation in Obsessive-Compulsive Disorder. Biol Psychiatry. 2019 May 1;85(9):735–43.

Barbe MT, Reker P, Hamacher S, Franklin J, Kraus D, Dembek TA, et al. DBS of the PSA and the VIM in essential tremor. Neurology 2018, 91:e543–e550.

Basha D, Dostrovsky JO, Lopez Rios AL, Hodaie M, Lozano AM, Hutchison WD. Beta oscillatory neurons in the motor thalamus of movement disorder and pain patients. Exp Neurol 2014, 261:782–790.

Beaty RE, Kenett YN, Christensen AP, Rosenberg MD, Benedek M, Chen Q, et al. Robust prediction of individual creative ability from brain functional connectivity. Proc Natl Acad Sci 2018 115:1087–1092.

Benabid AL, Benazzouz A, Gao D, Hoffmann D, Limousin P, Koudsie A, et al. Chronic Electrical Stimulation of the Ventralis Intermedius Nucleus of the Thalamus and of Other Nuclei as a Treatment for Parkinson’s Disease. Tech Neurosurg 1999, 5:5–30.

Benabid AL, Pollak P, Hoffmann D, Gervason C, Hommel M, Perret JE, et al. Long-term suppression of tremor by chronic stimulation of the ventral intermediate thalamic nucleus. Lancet 1991, 337:403–6.

Bhatia KP, Bain P, Bajaj N, Elble RJ, Hallett M, Louis ED, et al. Consensus Statement on the classification of tremors. from the task force on tremor of the International Parkinson and Movement Disorder Society. Mov Disord 2018, 33:75–87.

Boutet A, Ranjan M, Zohng J, Germann J, Xu D, Schwart ML, et al. Focused ultrasound thalamotomy location determines clinical benefits in patients with essential tremor. Brain 2018, 141, 3405, 3414.

Buckner RL, Krienen FM, Castellanos A, Diaz JC, Yeo BTT. The organization of the human cerebellum estimated by intrinsic functional connectivity. J Neurophysiol 2011, 106:2322–2345.

Cao H, Chén OY, Chung Y, Forsyth JK, McEwen SC, Gee DG, et al. Cerebello-thalamo-cortical hyperconnectivity as a state-independent functional neural signature for psychosis prediction and characterization. Nat Commun 2018, 9(1):3836.

Ceballos-Baumann AO, Boecker H, Fogel W, Alesch F, Bartenstein P, Conrad B, et al. Thalamic stimulation for essential tremor activates motor and deactivates vestibular cortex. Neurology 2001, 56:1347–1354.

Coenen VA, Allert N, Mädler B. A role of diffusion tensor imaging fiber tracking in deep brain stimulation surgery: DBS of the dentato-rubro-thalamic tract (drt) for the treatment of therapy-refractory tremor. Acta Neurochir (Wien) 2011a, 153:1579–1585.

Coenen VA, Allert N, Paus S, Kronenbürger M, Urbach H, Mädler B. Modulation of the Cerebello-Thalamo-Cortical Network in Thalamic Deep Brain Stimulation for Tremor. Neurosurgery 2014, 75:657–670.

Coenen VA, Mädler B, Schiffbauer H, Urbach H, Allert N. Individual Fiber Anatomy of the Subthalamic Region Revealed With Diffusion Tensor Imaging: A Concept to Identify the Deep Brain Stimulation Target for Tremor Suppression. Neurosurgery 2011b, 68:1069– 1076.

Coenen VA, Varkuti B, Parpaley Y, Skodda S, Prokop T, Urbach H, et al. Postoperative neuroimaging analysis of DRT deep brain stimulation revision surgery for complicated essential tremor. Acta Neurochir (Wien) 2017, 159(5):779–787.

Colebatch J, Findley LJ, Frackowiak RS, Marsden CD, Brooks DJ. Preliminary report: activation of the cerebellum in essential tremor. Lancet 1990. 336:1028–1030.

Cury RG, Fraix V, Castrioto A, Pérez Fernández MA, Krack P, Chabardes S, et al. Thalamic deep brain stimulation for tremor in Parkinson disease, essential tremor, and dystonia. Neurology 2017, 89:1416–1423.

Czarnecki K, Jones DT, Burnett MS, Mullan B, Matsumoto JY. SPECT perfusion patterns distinguish psychogenic from essential tremor. Parkinsonism Relat Disord 2011, 17:328– 332.

Darby RR, Horn A, Cushman F, Fox MD. Lesion network localization of criminal behavior. Proc Natl Acad Sci 2018, 115:601–6.

Deuschl G, Wenzelburger R, Löffler K, Raethjen J, Stolze H. Essential tremor and cerebellar dysfunction Clinical and kinematic analysis of intention tremor. Brain 2000, 123:1568– 1580.

Deuschl G, Raethjen J, Hellriegel H, Elble R. Treatment of patients with essential tremor. [Review]. Lancet Neurol 2011, 10:148–161.

Diamond A, Shahed J, Jankovic J. The effects of subthalamic nucleus deep brain stimulation on parkinsonian tremor. J Neurol Sci 2007, 260:199–203.

Eisinger RS, Wong J, Almeida L, Ramirez-Zamora A, Cagle JN, Giugni JC, et al. Ventral Intermediate Nucleus Versus Zona Incerta Region Deep Brain Stimulation in Essential Tremor. Mov Disord Clin Pract 2018, 5:75–82.

Fahn S, Tolosa E, Conceppcion M. Clinical rating scale for tremor In: Jankovic J, editor;, Tolosa E, editor., eds. Parkinson’s Disease and Movement Disorders. Baltimore, MD: Williams and Wilkins; 1993:271–280.

Fang W, Chen H, Wang H, Zhang H, Puneet M, Liu M, et al. Essential tremor is associated with disruption of functional connectivity in the ventral intermediate Nucleus-Motor Cortex-Cerebellum circuit. Hum Brain Mapp 2016, 37:165–178.

Fang W, Lv F, Luo T, Cheng O, Liao W, Sheng K, et al. Abnormal Regional Homogeneity in Patients with Essential Tremor Revealed by Resting-State Functional MRI. PLoS One 2013, 8(7):e69199.

Fenoy AJ, Schiess MC. Comparison of Tractography-Assisted to Atlas-Based Targeting for Deep Brain Stimulation in Essential Tremor. Mov Disord 2018, 00:1–7.

Fillard P, Descoteaux M, Goh A, Gouttard S, Jeurissen B, Malcolm J, et al. Quantitative evaluation of 10 tractography algorithms on a realistic diffusion MR phantom. Neuroimage 2011, 56:220–234.

Flora E Della, Perera CL, Cameron AL, Maddern GJ. Deep brain stimulation for essential tremor: A systematic review. Mov Disord 2010, 25:1550–9.

Foote KD, Seignourel P, Fernandez HH, Romrell J, Whidden E, Jacobson C, et al. Dual electrode thalamic deep brain stimulation for the treatment of posttraumatic and multiple sclerosis tremor. Neurosurgery 2006, 58:280–5.

Fytagoridis A, Blomstedt P. Complications and Side Effects of Deep Brain Stimulation in the Posterior Subthalamic Area. Stereotact Funct Neurosurg 2010, 88:88–93.

Fytagoridis A, Sandvik U, Åström M, Bergenheim T, Blomstedt P. Long term follow-up of deep brain stimulation of the caudal zona incerta for essential tremor. J Neurol Neurosurg Psychiatry 2012, 83:258–262.

Gallay MN, Jeanmonod D, Liu J, Morel A. Human pallidothalamic and cerebellothalamic tracts: anatomical basis for functional stereotactic neurosurgery. Brain Struct Funct 2008, 212:443–463.

Gibson WS, Jo HJ, Testini P, Cho S, Felmlee JP, Welker KM, et al. Functional correlates of the therapeutic and adverse effects evoked by thalamic stimulation for essential tremor. Brain 2016, 139:2198–2210.

Haber SN, Calzavara R. The cortico-basal ganglia integrative network: The role of the thalamus. Brain Res Bull 2009, 16;78(2-3):69–74.

Hamel W, Herzog J, Kopper F, Pinsker M, Weinert D, Müller D, et al. Deep brain stimulation in the subthalamic area is more effective than nucleus ventralis intermedius stimulation for bilateral intention tremor. Acta Neurochir (Wien) 2007, 149:749–758.

Hellwig B, Häußler S, Schelter B, Lauk M, Guschlbauer B, Timmer J, et al. Tremor-correlated cortical activity in essential tremor. Lancet 2001, 357:519–523.

Helmich RC, Toni I, Deuschl G, Bloem BR. The Pathophysiology of Essential Tremor and Parkinson’s Tremor. [Review]. Curr Neurol Neurosci Rep 2013, 13:378.

Herzog J, Fietzek U, Hamel W, Morsnowski A, Steigerwald F, Schrader B, et al. Most effective stimulation site in subthalamic deep brain stimulation for Parkinson’s disease. Mov Disord 2004, 19:1050–4.

Horn A. A structural group-connectome in standard stereotactic (MNI) space. Data Br 2015, 5:292–6.

Horn A, Blankenburg F. Toward a standardized structural-functional group connectome in MNI space. Neuroimage 2016, 124:310–322.

Horn A, Kühn AA, Merkl A, Shih L, Alterman R, Fox M. Probabilistic conversion of neurosurgical DBS electrode coordinates into MNI space. Neuroimage 2017a, 150:395– 404.

Horn A, Ostwald D, Reisert M, Blankenburg F. The structural–functional connectome and the default mode network of the human brain. Neuroimage 2014, 102:142–151.

Horn A, Reich M, Vorwerk J, Li N, Wenzel G, Fang Q, etl. Connectivity Predicts deep brain stimulation outcome in Parkinson disease. Ann Neurol 2017b, 82:67–78.

Hoskovcová M, Ulmanová O, Šprdlík O, Sieger T, Nováková J, Jech R, et al. Disorders of Balance and Gait in Essential Tremor Are Associated with Midline Tremor and Age. The Cerebellum 2013, 12:27–34.

Jenkins IH, Bain PG, Colebatch JG, Thompson PD, Findley LJ, Frackowiak RSJ, et al. A positron emission tomography study of essential tremor: Evidence for overactivity of cerebellar connections. Ann Neurol 1993, 34:82–90.

Joutsa J, Horn A, Hsu J, Fox MD. Localizing parkinsonism based on focal brain lesions. Brain 2018a, 141:2445–2456.

Joutsa J, Shih LC, Horn A, Reich MM, Wu O, Rost NS, et al. Identifying therapeutic targets from spontaneous beneficial brain lesions. Ann Neurol 2018b, 84:153–7.

Kilbane C, Ramirez-Zamora A, Ryapolova-Webb E, Qasim S, Glass GA, Starr PA, et al.Pallidal stimulation for Holmes tremor: clinical outcomes and single-unit recordings in 4 cases. J Neurosurg 2015, 122:1306–1314.

Klein JC, Barbe MT, Seifried C, Baudrexel S, Runge M, Maarouf M, et al. The tremor network targeted by successful VIM deep brain stimulation in humans. Neurology 2012, 78:787– 795.

Kumar R, Lozano AM, Sime E, Lang AE. Long-term follow-up of thalamic deep brain stimulation for essential and parkinsonian tremor. Neurology 2003, 61:1601–4.

Mandat T, Koziara H, Tutaj M, Rola R, Bonicki W, Nauman P. Thalamic deep brain stimulation for tremor among multiple sclerosis patients. Neurol Neurochir Pol 2010, 44:542–5.

Marsden JF, Ashby P, Limousin-Dowsey P, Rothwell JC, Brown P. Coherence between cerebellar thalamus, cortex and muscle in man: cerebellar thalamus interactions. Brain 2000, 123:1459–1470.

McAuley JH, Marsden CD. Physiological and pathological tremors and rhythmic central motor control. [Review]. Brain 2000, 123:1545–1567.

McFarland NR, Haber SN. Thalamic relay nuclei of the basal ganglia form both reciprocal and nonreciprocal cortical connections, linking multiple frontal cortical areas. [Review]. J Neurosci 2002, 22:8117–8132.

Middlebrooks EH, Tuna IS, Almeida L, Grewal SS, Wong J, Heckman MG, et al. Structural connectivity–based segmentation of the thalamus and prediction of tremor improvement following thalamic deep brain stimulation of the ventral intermediate nucleus. NeuroImage Clin 2018, 20:1266–1273.

Milosevic L, Kalia SK, Hodaie M, Lozano AM, Popovic MR, Hutchison WD. Physiological mechanisms of thalamic ventral intermediate nucleus stimulation for tremor suppression. Brain 2018, 141:2142–2155.

Mueller K, Jech R, Hoskovcová M, Ulmanová O, Urgošík D, Vymazal J,et al. General and selective brain connectivity alterations in essential tremor: A resting state fMRI study. NeuroImage Clin 2017, 16:468–476.

Obwegeser AA, Uitti RJ, Turk MF, Strongosky AJ, Wharen RE. Thalamic stimulation for the treatment of midline tremors in essential tremor patients. Neurology 2000, 54:2342–4.

Pahwa R, Lyons KE, Wilkinson SB, Simpson RK, Ondo WG, Tarsy D, et al. Long-term evaluation of deep brain stimulation of the thalamus. J Neurosurg 2006, 104:506–512.

Papavassiliou E, Rau G, Heath S, Abosch A, Barbaro NM, Larson PS, et al. Thalamic deep brain stimulation for essential tremor: relation of lead location to outcome. Neurosurgery 2004, 54(5):1120–29.

Pedrosa DJ, Reck C, Florin E, Pauls KAM, Maarouf M, Wojtecki L, et al. Essential tremor and tremor in Parkinson’s disease are associated with distinct ‘tremor clusters’ in the ventral thalamus. Exp Neurol 2012, 237:435–443.

Penfield W, Boldrey E. Somatic motor and sensory representation in the cerebral cortex of man as studied by electrical stimulation. Brain 1937, 60:389–443.

Pinto AD, Lang AE, Chen R. The cerebellothalamocortical pathway in essential tremor. Neurology 2003, 60:1985–7.

Plaha P, Javed S, Agombar D, O’ Farrell G, Khan S, Whone A, et al. Bilateral caudal zona incerta nucleus stimulation for essential tremor: outcome and quality of life. J Neurol Neurosurg Psychiatry 2011, 82:899–904.

Plaha P, Khan S, Gill SS. Bilateral stimulation of the caudal zona incerta nucleus for tremor control. J Neurol Neurosurg Psychiatry 2008, 79:504–513.

Plaha P, Patel NK, Gill SS. Stimulation of the subthalamic region for essential tremor. J Neurosurg 2004, 101:48–54.

Pouratian N, Zheng Z, Bari AA, Behnke E, Elias WJ, Desalles AAF. Multi-institutional evaluation of deep brain stimulation targeting using probabilistic connectivity-based thalamic segmentation. J Neurosurg 2011, 115:995–1004.

Putzke JD, Uitti RJ, Obwegeser AA, Wszolek ZK, Wharen RE. Bilateral thalamic deep brain stimulation: midline tremor control. J Neurol Neurosurg Psychiatry 2005, 76:684–690.

Raethjen J, Deuschl G. The oscillating central network of Essential tremor. [Review]. Clin Neurophysiol 2012, 123:61–4.

Reich MM, Brumberg J, Pozzi NG, Marotta G, Roothans J, Åström M, et al. Progressive gait ataxia following deep brain stimulation for essential tremor: adverse effect or lack of efficacy? Brain 2016 Nov 1;139(11):2948–56.

Reisert M, Mader I, Anastasopoulos C, Weigel M, Schnell S, Kiselev V. Global fiber reconstruction becomes practical. Neuroimage 2011, 54(2):955–62.

Sammartino F, Krishna V, King NKK, Lozano AM, Schwartz ML, Huang Y, et al. Tractography-Based Ventral Intermediate Nucleus Targeting: Novel Methodology and Intraoperative Validation. Mov Disord 2016, 31:1217–1225.

Sandvik U, Koskinen L-O, Lundquist A, Blomstedt P. Thalamic and Subthalamic Deep Brain Stimulation for Essential Tremor. Neurosurgery 2012, 70:840–6.

Schnitzler A, Münks C, Butz M, Timmermann L, Gross J. Synchronized brain network associated with essential tremor as revealed by magnetoencephalography. Mov Disord 2009, 24:1629–1635.

Sharifi S, Nederveen AJ, Booij J, van Rootselaar A-F. Neuroimaging essentials in essential tremor: a systematic review. [Review]. NeuroImage Clin 2014, 5:217–231.

Tasker RR. Deep brain stimulation is preferable to thalamotomy for tremor suppression. Surg Neurol 1998, 49:145–154.

Tuleasca C, Najdenovska E, Régis J, Witjas T, Girard N, Champoudry J, et al. Pretherapeutic functional neuroimaging predicts tremor arrest after thalamotomy. Acta Neurol Scand 2018a, 137:500–8.

Tuleasca C, Najdenovska E, Régis J, Witjas T, Girard N, Champoudry J, et al. Clinical response to Vim’s thalamic stereotactic radiosurgery for essential tremor is associated with distinctive functional connectivity patterns. Acta Neurochir (Wien) 2018b, 160:611–624.

Tuleasca C, Witjas T, Najdenovska E, Verger A, Girard N, Champoudry J, et al. Assessing the clinical outcome of Vim radiosurgery with voxel-based morphometry: visual areas are linked with tremor arrest! Acta Neurochir (Wien) 2017, 159:2139–2144.

Tuleasca C, Witjas T, Van de Ville D, Najdenovska E, Verger A, Girard N, et al. Right Brodmann area 18 predicts tremor arrest after Vim radiosurgery: a voxel-based morphometry study. Acta Neurochir (Wien) 2018c, 160:603–9.

Vayssiere N, van der Gaag N, Cif L, Hemm S, Verdier R, Frerebeau P, et al. Deep brain stimulation for dystonia confirming a somatotopic organization in the globus pallidus internus. J Neurosurg 2004, 101(2):181–8.

Weigand A, Horn A, Caballero R, Cooke D, Stern AP, Taylor SF, et al. Prospective Validation That Subgenual Connectivity Predicts Antidepressant Efficacy of Transcranial Magnetic Stimulation Sites. Biol Psychiatry 2018, 84:28–37.

Wills AJ, Jenkins LH, Thompson PD, Findley LJ, Brooks DJ. A Positron Emission Tomography Study of Cerebral Activation Associated With Essential and Writing Tremor. Arch Neurol 1995, 52:299–305.

Yamada K, Akazawa K, Yuen S, Goto M, Matsushima S, Takahata A, et al. MR Imaging of Ventral Thalamic Nuclei. Am J Neuroradiol 2010, 31:732–5.

Zhang K, Bhatia S, Oh MY, Cohen D, Angle C, Whiting D. Long-term results of thalamic deep brain stimulation for essential tremor. J Neurosurg 2010, 112:1271–6.

