## Supplementary material for "Connectivity profile of thalamic deep brain stimulation to effectively treat essential tremor": Fig. S1, Fig. S2

### Representative Single-Subject Prospective Validation

#### Imaging Acquisition and Pre-processing

High-resolution anatomical images were obtained using a standard magnetization-prepared rapid gradient-echo (MPRAGE) sequence (TR = 2300 ms, TE = 2.32 ms, isotropic voxel-size of 0.9 × 0.9 × 0.9 $mm^3$, 192 slices). Diffusion-weighted images (DTI) were obtained using 3.0-Tesla Siemens Magnetom SKYRA (Siemens Medical System, Erlangen, Germany) (voxel-size of 2 × 2 × 2 $mm^3$, 60 slices). An effective b-value of 1000 s/$mm^2$ was used for each of the 40 diffusion encoding directions. In addition, 1 volume without diffusion weighting (b-value = 0 s/$mm^2$) equally distributed throughout the scan were acquired (b0-images). Motion and distortion of DTI sequences were first corrected by using FSL eddy_correct function. DTI sequences were then co-registered to anatomical images (MPRAGE) using SPM implemented in Lead-DBS.

#### Connectivity Estimation

A dMRI diffusion scheme implemented in DSI-Studio (implemented in Lead-DBS) was used. Specifically, diffusion data were reconstructed using generalized q-sampling imaging (Yeh *et al.* 2011) with a diffusion sampling length ratio of 1.25. The restricted diffusion was quantified using restricted diffusion processing (Yeh *et al.* 2013). A deterministic fiber tracking algorithm (Yeh *et al.* 2013) was used, the angular threshold was 60 degrees. The step size was 0.86 mm. The anisotropy threshold was determined automatically by DSI Studio. Tracks with length below 10 mm were discarded. A total of 20000 tracts were calculated per subject. The whole- brain fiber set was then normalized into standard-stereotactic space following the approach described in (Horn *et al*. 2017a, Horn *et al.* 2017b) as implemented in Lead-DBS.

#### References

1. Yeh FC, Tseng WYI: NTU-90: A high angular resolution brain atlas constructed by q-space diffeomorphic reconstruction. Neuroimage, 2011. 58: 91–99.
2. Yeh FC, Verstynen TD, Wang Y, Fernández-Miranda JC, Tseng WYI: Deterministic diffusion fiber tracking improved by quantitative anisotropy. PLoS One 2013. 15;8(11):e80713.
3. Horn A, Neumann W-J, Degen K, Schneider G-H, Kühn AA: Toward an electrophysiological “sweet spot” for deep brain stimulation in the subthalamic nucleus. Hum Brain Mapp, 2017. 38(7):3377-90.
4. Horn A, Ostwald D, Reisert M, Blankenburg F: The structural-functional connectome and the default mode network of the human brain. Neuroimage, 2014. 102: 142–51.

Supplementary Table S1: Tremor score and demographic characteristics of study cohort

| ***Patient #*** | ***Age*** | ***Sex*** | ***Age at Diagnosis*** | ***Disease Duration*** | ***Total FTM score % improvement*** | ***Right upper limb FTM subscore % improvement*** | ***Left upper limb FTM subscore % improvement*** | ***Bilateral upper limb FTM subscore % improvement*** | ***Head FTM subscore % improvement*** |
|---|---|---|---|---|---|---|---|---|---|
| ***#1*** | 74 | M | 51 | 23 | 95.5% | 92.9% | 100.0% | 96.2% | 66.7% |
| ***#2*** | 52 | F | 21 | 33 | 59.5% | 76.9% | 71.4% | 75.0% | 14.3% |
| ***#3*** | 85 | M | 62 | 16 | 46.5% | 33.3% | 41.7% | 37.5% | 75.0% |
| ***#4*** | 73 | F | 47 | 20 | 48.8% | 62.5% | 30.8% | 48.3% | 100.0% |
| ***#5*** | 77 | M | 65 | 12 | 78.0% | 71.4% | 91.7% | 80.8% | 83.3% |
| ***#6*** | 79 | F | 40 | 44 | 33.3% | 16.7% | 27.3% | 23.5% | NaN |
| ***#7*** | 75 | F | 57 | 8 | 58.6% | 62.5% | 33.3% | 47.1% | 100.0% |
| ***#8*** | 59 | F | 44 | 15 | 25.0% | 0.0% | 37.5% | 25.0% | NaN |
| ***#9*** | 78 | F | 64 | 14 | 90.6% | 100.0% | 76.9% | 88.5% | 100.0% |
| ***#10*** | 74 | M | 38 | 40 | 41.3% | 43.8% | 33.3% | 39.3% | 40.0% |
| ***#11*** | 59 | F | 13 | 43 | 88.0% | 80.0% | 100.0% | 90.0% | NaN |
| ***#12*** | 69 | F | 35 | 32 | 60.0% | 20.0% | 71.4% | 57.9% | NaN |
| ***#13*** | 30 | M | 15 | 3 | 18.8% | 20.0% | 42.9% | 33.3% | 0.0% |
| ***#14*** | 70 | M | 51 | 11 | 92.6% | 83.3% | 90.0% | 87.5% | 100.0% |
| ***#15*** | 78 | F | 12 | 48 | 75.0% | 69.2% | 72.7% | 70.8% | 87.5% |
| ***#16*** | 78 | F | 53 | 13 | 76.2% | 45.5% | 76.9% | 62.5% | 100.0% |
| ***#17*** | 82 | F | 65 | 11 | 76.3% | 66.7% | 86.7% | 77.8% | 100.0% |
| ***#18*** | 84 | F | 66 | 10 | 26.9% | 10.0% | 33.3% | 21.1% | NaN |
| ***#19*** | 86 | F | 53 | 25 | 75.7% | 62.5% | 90.0% | 73.1% | 100.0% |
| ***#20*** | 79 | F | 50 | 20 | 70.0% | 78.6% | 50.0% | 64.3% | NaN |
| ***#21*** | 79 | F | 51 | 22 | 69.0% | 70.0% | 70.0% | 70.0% | 100.0% |
| ***#22*** | 83 | F | 40 | 38 | 61.8% | 40.0% | 61.5% | 52.2% | NaN |
| ***#23*** | 74 | M | 15 | 54 | 63.2% | 75.0% | 42.9% | 60.0% | NaN |
| ***#24*** | 84 | M | 35 | 40 | 56.7% | 87.5% | 40.0% | 61.1% | 80.0% |
| ***#25*** | 77 | M | 60 | 9 | 37.9% | 50.0% | 25.0% | 36.4% | NaN |
| ***#26*** | 83 | F | 58 | 15 | 81.6% | 90.9% | 66.7% | 76.9% | NaN |
| ***#27*** | 90 | M | 66 | 15 | 56.8% | 60.0% | 50.0% | 53.8% | 50.0% |
| ***#28*** | 79 | F | 15 | 64 | 84.6% | 78.6% | 84.6% | 81.5% | 100.0% |
| ***#29*** | 72 | M | 65 | 15 | 90.2% | 93.8% | 84.6% | 89.7% | 80.0% |
| ***#30*** | 72 | F | 33 | 30 | 48.6% | 9.1% | 73.3% | 46.2% | NaN |
| ***#31*** | 55 | F | 14 | 30 | 35.0% | 33.3% | 12.5% | 21.4% | 100.0% |
| ***#32*** | 93 | F | 45 | 20 | 78.8% | 57.1% | 75.0% | 68.4% | 100.0% |
| ***#33*** | 76 | M | 59 | 13 | 52.4% | 33.3% | 62.5% | 50.0% | 100.0% |
| ***#34*** | 80 | F | 70 | 10 | 67.6% | 75.0% | 50.0% | 69.6% | NaN |
| ***#35*** | 61 | M | 46 | 15 | 59.5% | 71.4% | 47.4% | 57.6% | NaN |
| ***#36*** | 75 | F | 30 | 45 | 72.5% | %83.3 | 61.5% | 74.2% | NaN |
| ***Mean*** | 74.48 | | 44.18 | 24.42 | 62.2% | 56.8% | 60.8% | 59.6% | 80.8% |
| ***SD*** | 12.19 | | 18.12 | 14.94 | 21.2% | 27.7% | 24.4% | 21.9% | 29.5% |

FTM – Fahn-Tolosa-Marin tremor rating scale, M – male, F – female. NaN – not a number, indicates 0 tremor subscore at baseline and postoperatively. Head tremor subscore consisted of FTM subscore items for head, tongue, voice, speaking and face tremor.

### Supplementary Table S2: Individual patients clinical DBS settings

| ***Patient #*** | ***DBS electrode model*** | ***Right electrode settings*** | ***Amplitude (V/mA)*** | ***Frequency (Hz)*** | ***Pulse Width (μs)*** | ***Left electrode settings*** | ***Amplitude (V/mA)*** | ***Frequency (Hz)*** | ***Pulse Width (μs)*** |
|---|---|---|---|---|---|---|---|---|---|
| ***#1*** | Medtronic 3387 | 1 -ve, case +ve | 1.5V | 130 | 60 | 4 -ve, case +ve | 2.5V | 130 | 60 |
| ***#2*** | Medtronic 3387 | 0 -ve, case +ve | 1.9V | 110 | 60 | 4 -ve, case +ve | 2.4V | 110 | 60 |
| ***#3*** | Medtronic 3387 | 1 -ve, case +ve | 2.1V | 130 | 60 | 5 -ve, case +ve | 2V | 130 | 60 |
| ***#4*** | Medtronic 3387 | 1 -ve, case +ve | 2V | 130 | 60 | 5 -ve, case +ve | 1.5V | 130 | 60 |
| ***#5*** | Medtronic 3387 | 1 -ve, case +ve | 2.4V | 160 | 60 | 5 -ve, case +ve | 3.6V | 160 | 60 |
| ***#6*** | Medtronic 3387 | 1 -ve, case +ve | 2.8V | 130 | 60 | 5 -ve, case +ve | 2.8V | 130 | 60 |
| ***#7*** | Medtronic 3387 | 1 -ve, case +ve | 2V | 180 | 60 | 5 -ve, case +ve | 3.1V | 180 | 60 |
| ***#8*** | Medtronic 3387 | 2 -ve, case +ve | 1.6V | 130 | 60 | 6 -ve, case +ve | 2.4V | 130 | 60 |
| ***#9*** | St. Jude ActiveTip (6142-6145) | 0 -ve, case +ve | 2.5mA | 130 | 100 | 4 -ve, case +ve | 5.5mA | 130 | 75 |
| ***#10*** | Medtronic 3387 | 0 and 1 -ve, case +ve | 2.7V , 1.3V | 120 | 60 | 8 and 9 -ve, case +ve | 0.9V , 0.8V | 120 | 60 |
| ***#11*** | Boston Scientific Vercise Directed | 13 and 14 -ve, case +ve | 3mA | 130 | 60 | 4 and 7 -ve, case +ve | 3mA | 130 | 60 |
| ***#12*** | Medtronic 3387 | 1 -ve, case +ve | 2.4V | 130 | 60 | 9 -ve, case +ve | 1.7 | 130 | 6 |
| ***#13*** | Medtronic 3387 | 1 -ve, case +ve | 1.5V | 130 | 90 | 8 -ve, case +ve | 2.5 | 130 | 90 |
| ***#14*** | Medtronic 3387 | 1 -ve, case +ve | 3V | 130 | 60 | 5 -ve, case +ve | 2V | 130 | 60 |
| ***#15*** | Medtronic 3387 | 1 and 2, case +ve | 1.8V | 180 | 60 | 5 -ve, case +ve | 3V | 180 | 60 |
| ***#16*** | Medtronic 3387 | 1 -ve, case +ve | 2.8V | 210 | 90 | 5 -ve, case +ve | 2.7V | 210 | 90 |
| ***#17*** | Medtronic 3387 | 1 -ve, case +ve | 1.7V | 130 | 60 | 9 -ve, case +ve | 1.7V | 130 | 60 |
| ***#18*** | Medtronic 3387 | 2 -ve, case +ve | 2.5V | 125 | 60 | 4 and 5 -ve, case +ve | 2V , 2.5V | 125 | 60 |
| ***#19*** | Medtronic 3387 | 3 -ve, case +ve | 3V | 155 | 90 | 5 and 6 -ve, case +ve | 4.5V | 167 | 60 |
| ***#20*** | Medtronic 3387 | 1 -ve, case +ve | 3.1V | 130 | 60 | 7 -ve, case +ve | 1.4V | 130 | 60 |
| ***#21*** | Medtronic 3387 | 1 -ve, case +ve | 2V | 130 | 60 | 9 -ve, case +ve | 2.3V | 130 | 60 |
| ***#22*** | Medtronic 3387 | 0 -ve, case +ve | 2V | 130 | 60 | 4 -ve, case +ve | 2V | 130 | 60 |
| ***#23*** | Medtronic 3387 | 1 -ve, case +ve | 2.4V | 130 | 60 | 9 -ve, case +ve | 2.9V | 130 | 150 |
| ***#24*** | Medtronic 3387 | 1 -ve, case +ve | 2.3V | 130 | 60 | 5 -ve, case +ve | 2V | 130 | 60 |
| ***#25*** | Medtronic 3387 | 1 and 2 -ve, case +ve | 3.3V , 3V | 125 | 90 | 5 and 6 -ve, case +ve | 3.3V | 125 | 90 |
| ***#26*** | Medtronic 3387 | 1 -ve, case +ve | 3V | 130 | 60 | 5 -ve, case +ve | 3V | 130 | 60 |
| ***#27*** | Medtronic 3387 | 1-ve, case +ve | 2.3V | 130 | 60 | 5 -ve, case +ve | 2V | 130 | 60 |
| ***#28*** | Medtronic 3387 | 1-ve, case +ve | 4.5V | 130 | 60 | 4 -ve, case +ve | 3V | 130 | 60 |
| ***#29*** | Medtronic 3387 | 2-ve, case +ve | 2.8V | 130 | 60 | 9 -ve, case +ve | 2.6V | 130 | 60 |
| ***#30*** | Medtronic 3387 | 1-ve, case +ve | 3V | 210 | 60 | 5 and 6 -ve, case +ve | 5.5 | 210 | 60 |
| ***#31*** | Medtronic 3387 | 2-ve, case +ve | 3V | 180 | 60 | 6 -ve, case +ve | 2.5 | 180 | 60 |
| ***#32*** | Medtronic 3387 | 1-ve, case +ve | 3.4V | 130 | 90 | 9 -ve, case +ve | 2.7V | 130 | 90 |
| ***#33*** | Medtronic 3387 | 0-ve, case +ve | 2V | 150 | 60 | 10 -ve, case +ve | 3V | 150 | 90 |
| ***#34*** | Medtronic 3387 | 3 and 4 -ve, case +ve | 2.1V , 4.1V | 125 | 60 | 7 -ve, case +ve | 2V | 125 | 60 |
| ***#35*** | Boston Scientific Vercise Directed | 11 and 12 -ve, case +ve | 2.4mA | 174 | 80 | 3 and 4 -ve, case +ve | 2.8mA | 174 | 80 |
| ***#36*** | Medtronic 3387 | 0 -ve, 1 +ve | 2V | 180 | 60 | 8 -ve, 9 +ve | 2.3V | 180 | 60 |

μs - microsecond, V - volts, mA - milliampere, Hz – hertz

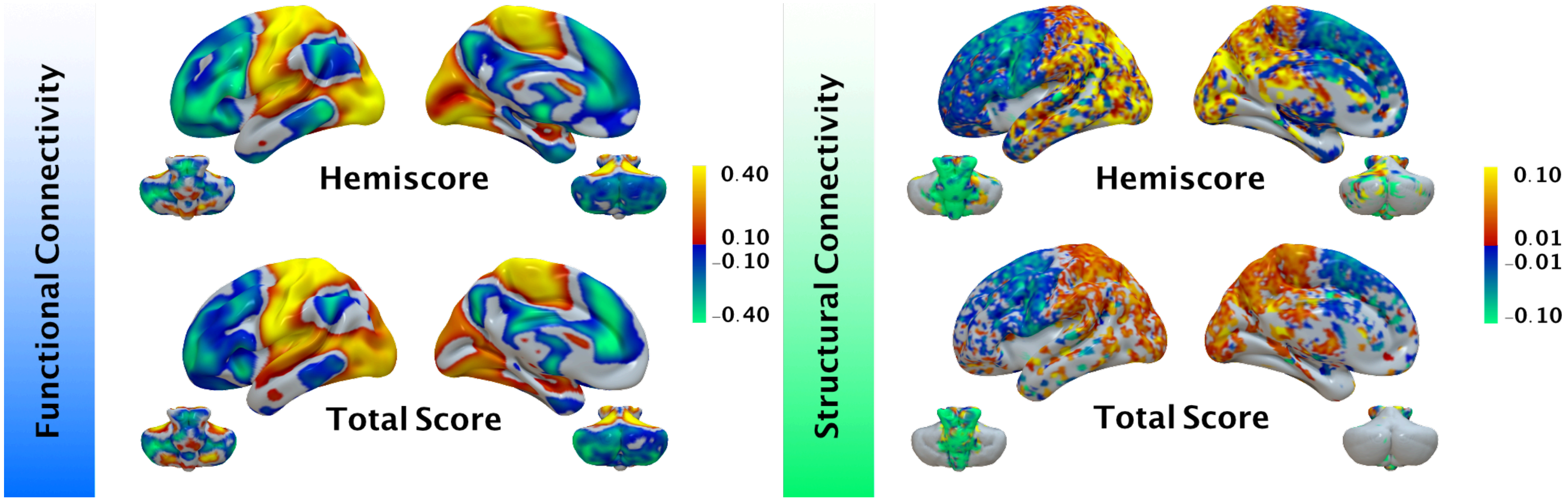


Figure S1: Direct comparison between hand-tremor analysis (that formed the main part of the analysis, top) and improvements on full tremor scores (bottom).

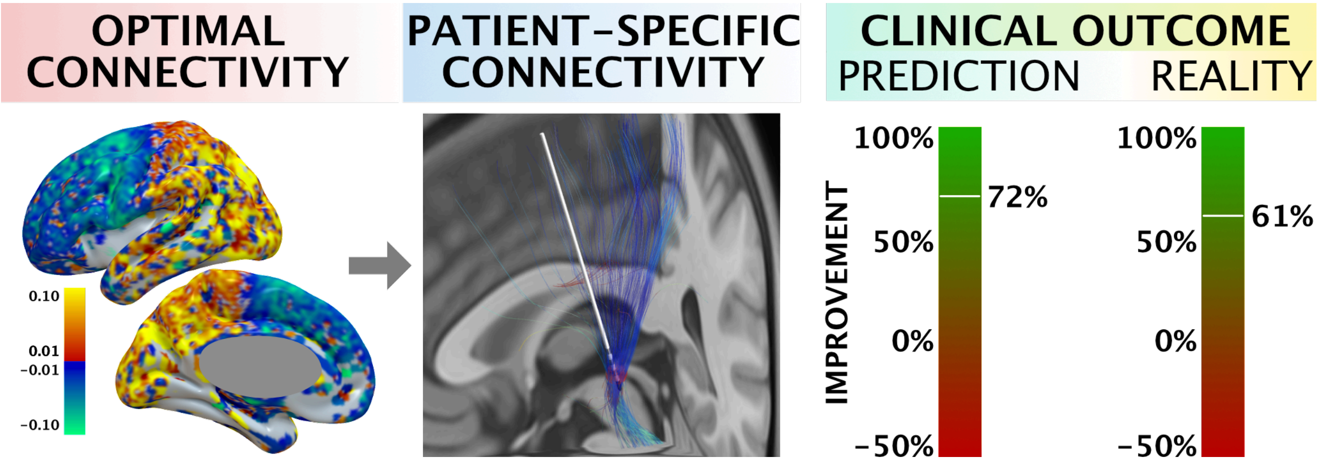


Figure S2: Patient-specific connectivity (middle) was used to predict a prospective patient upper limb improvement based on normative connectome structural-model (left). Predicted and empirical improvements are scaled in left panel.
